# Moving through everyday life: distinct developmental timescales for head-rotation magnitude and rotation efficiency in infancy

**DOI:** 10.64898/2026.08.09.742392

**Authors:** Zachary J. Petroff, Rajat Kapgate, T. Rowan Candy, Linda B. Smith, Kathryn Bonnen

## Abstract

Infants actively shape their experience through self-generated movement. They need to move their head efficiently to explore and interact with their environment and to hold their head still to sustain attention. Controlled laboratory studies have documented the importance of head movements in orienting and stabilizing infant visual attention, but little is known about how head control develops as infants go about daily life, the setting in which development actually unfolds. We analyzed 383 hours of egocentric video collected from 88 infants aged 3 weeks to 29 months using head-mounted cameras in infants’ homes. Using a visual-odometry algorithm, we quantified rotational magnitude, the size of head movements; and rotational efficiency, the directional coherence of head movements. Head-rotation magnitude increased across the first year, and rotational efficiency improved throughout development.

## Introduction

An individual’s experience of the world depends on movements of their body and how they coordinate actions. These behaviors change systematically over the first two years of human infancy (Kretch et al., 2014; Smith et al., 2015; Adolph and Hoch, 2019; Franchak, 2020). Because head position and movements contribute significantly to the direction and stability of visual input, head control is central to the developmental coordination of visual experience. Indeed, the role of head control extends broadly in development: head movements and head stabilization have been implicated not only in developmental changes in visual experience (Kretch et al., 2014; Fausey et al., 2016; Franchak et al., 2024) and in learning from those experiences (Yu and Smith, 2012; Smith, 2013; Mendez et al., 2024), but also in the reorganization of sensory-motor systems with each new motor milestone (Hadders-Algra et al., 1996; Bril and Ledebt, 1998; Bertenthal and von Hofsten, 1998; Adolph and Franchak, 2017) and the development of attentional control and individual differences in attention (Friedman et al., 2005; Klingberg et al., 2002; Borjon et al., 2021).

A large literature indicates that head control progresses during the first year, from a limited ability to lift and maneuver the head, to the use of head orientation in guiding reaching, to a stabilized head in early walking, and a stabilized head in sustained attention and manual actions on objects (for reviews, see Adolph and Franchak (2017)). Studies of motor development also reveal that advances in other motor abilities can change biomechanical demands and disrupt infant head control, leading to periods of reorganization (Bril and Ledebt, 1998; Bertenthal and von Hofsten, 1998; Claxton et al., 2014; Adolph and Franchak, 2017). This occurs while head and postural control have also been linked to attentional control and the development of sustained attention during infancy (Borjon et al., 2021), with poor control identified as a risk factor for attentional disorders (Friedman et al., 2005; Klingberg et al., 2002). Together, these findings demonstrate that head control is not a single milestone but a developing capacity that supports a variety of developing infant behaviors.

Despite this convergent evidence, our current understanding of the development of infant head movements is based primarily on experiments in controlled laboratory environments, focused on specific skills at specific ages. What is missing is an account of how head control develops continuously across infancy, as infants move through the changing contexts of daily life. Here we quantify head movements of a cross-sectional sample of 88 infants from 3 weeks to 29 months of age, drawn from the dataset described in Fausey et al. (2016). We use head-mounted cameras and recent advances in visual odometry to characterize three aspects of infants’ head rotations during daily life at home: *net rotation*, the overall change in head orientation across short time windows; *total rotation*, the cumulative amount of head movement within those windows; and *efficiency*, the degree to which total movement contributes to a coherent change in orientation. By sampling head movements as infants go about everyday activity, we evaluate the development of head control within the natural setting in which infant development unfolds. Together, net rotation, total rotation, and efficiency capture how much infants move their head and how directly those movements are organized.

We find that head-rotation magnitude increases over the first year of life, whereas rotational efficiency continues to increase across the full age range sampled, through 29 months. This pattern suggests that different aspects of head control follow distinct developmental timelines. Early in development, infants produce increasingly larger head rotations, likely reflecting gains in strength, postural control, and opportunities for active exploration. Over a longer period, infants’ head movements also become more efficient, with a greater proportion of their total movement contributing to coherent changes in orientation.

## 2 Results

We characterized infant head rotations by estimating changes in the orientation of a head-mounted camera worn by infants (n=88, aged 3 wk to 29 mo; see Figure. 1). The data consisted of 383 hours of video. Because the sensory and motor capabilities of an infant change rapidly during the first two years of life, we sought to analyze the effects over time. To achieve this in our analysis, the recordings were grouped into ten 3-month age bins based on the age of each child at the time of recording.

**Figure 1:**
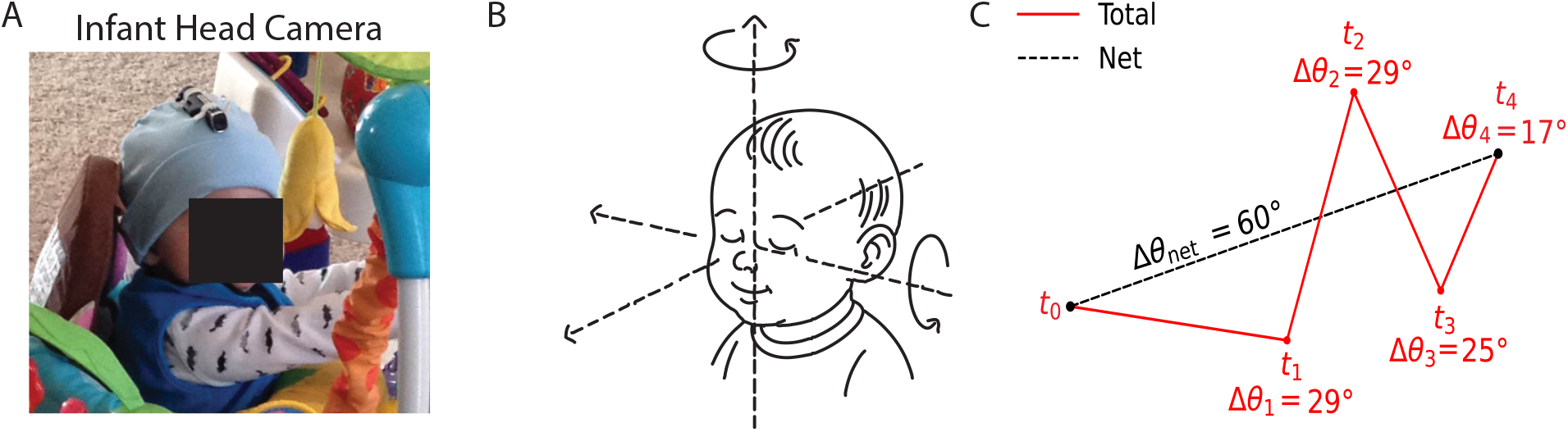
Measuring head rotation in the home environment. **(a) Recording egocentric video in the home environment.**Infants wore a lightweight Looxcie2 camera secured using a snug-fitting beanie during at-home activity. **(b) Head rotation estimation**. A visual odometry algorithm is used to approximate 3-dimensional head rotation. **(c) Two-dimensional projection of the head rotation sequence**. The rotation trajectory is projected into 2D to illustrate the distinction between total and net rotation. Total rotation is the cumulative angular change across successive orientations, whereas net rotation is the smallest angular difference between the initial and final orientations. Rotation efficiency is therefore computed as 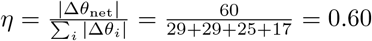.

**Figure 2.**
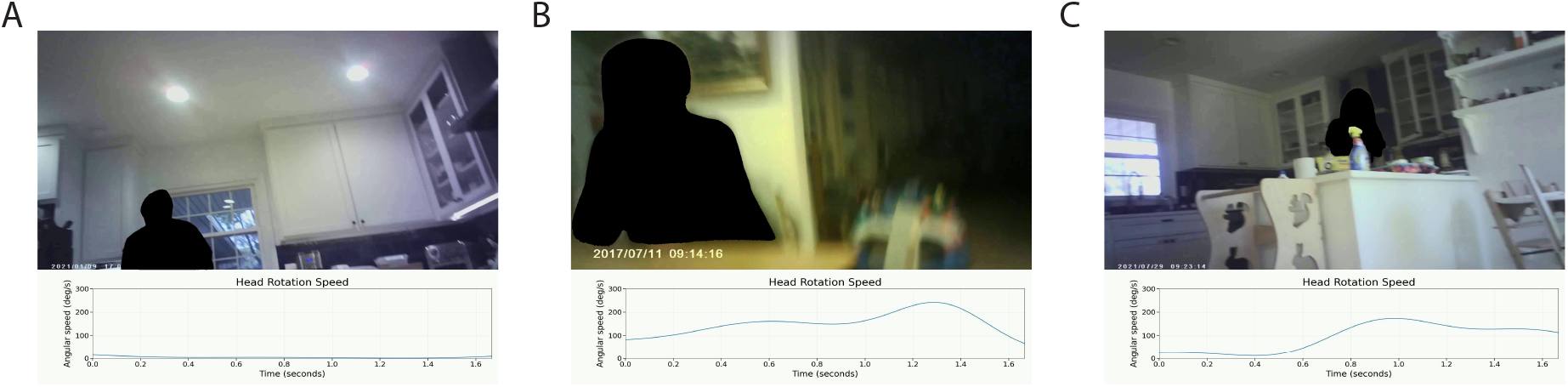
Examples of head rotation across infancy. **(a)** In the 3-month recording, head movement is limited and consists primarily of small stabilizing adjustments. **(b)** In the 6-month recording, head movement is more frequent and variable, with near-continuous motion accompanied by repeated stabilization. **(c)** In the 9-month recording, head movements are more organized, with a clearer distinction between periods of active reorientation and relative stillness. *Note*. Caregivers were masked in the example videos to protect their privacy while permitting the examples to be shared.

Notably, this method estimates head rotation relative to the environment. Thus we cannot distinguish between head rotations where the head rotates relative to the body from head rotations where the head rotates with the body.

To measure changes in head orientation using the egocentric video, we employed the method outlined in Yang and Ramanan (2021). This method was chosen because of its robustness in the presence of dynamic scene elements. This property is crucial for our dataset which features frequent close-range motion of hands, toys, and faces. The algorithm estimates rotation and translation between each frame pair. For these analyses we retained only the rotational component. Outliers and high-frequency noise were removed from the resulting rotation estimates using robust outlier rejection followed by temporal low-pass filtering. A more detailed description of this method and its validation can be found in the Materials and Methods section.

The rotation between two video frames is represented by a rotation matrix. To estimate head rotation over longer time intervals, we combined a series of these rotation matrices and computed two complementary measures of head rotation over time:

- **total rotation**: the cumulative amount of rotation that occurs within the time interval.
- **net rotation**: the most direct rotation between the start and end of a time interval.

We use these measurements of total rotation and net rotation to compute an efficiency metric for each movement interval (t=0.5 seconds): efficiency = net rotation / total rotation. When the net rotation is of similar value to the total rotation, the efficiency metric is high or close to one, indicating a perfectly directed movement. Conversely, if the total rotation is large but the net rotation is small, efficiency is low or close to zero, indicating movement that cancels itself out. This is common with very young babies who do not have strong neck control. The schematics in figure 1c illustrate efficient and inefficient head rotation trajectories for scenarios where the total head rotation is large or small.

### 2.1 Head rotation magnitudes increase during the first year of life

In general, we found that older infants move their head more and make more large head rotations than younger infants. Across the full age range, both total rotation and net rotation were greater in older infants (total: *β* = 0.026°/week, *SE* = 0.010, *z* = 2.634, *p* = 0.008; net: *β* = 0.021°/week, *SE* = 0.006, *z* = 3.547, *p <* 0.001). This effect was driven by the first 48 weeks of life (total: *β* = 0.148°/week, *SE* = 0.028, *z* = 5.278, *p <* 0.001; net: *β* = 0.099°/week, *SE* = 0.016, *z* = 6.008, *p <* 0.001); and no significant linear trend was observed after 48 weeks (total: *β* = *−*0.012, *SE* = 0.023, *z* = *−*0.536, *p* = 0.592; net: *β* = *−*0.001, *SE* = 0.014, *z* = *−* 0.084, *p* = 0.933). Figure 3a shows the distribution of total rotation across age groups, and Figure 4a shows the median total rotation per age group. Similarly, Figure 3b shows the distribution of net rotation, and Figure 4b shows the median net rotation per age group. Thus, both total rotation and net rotation increase over the first year of life, and subsequently fluctuate.

**Figure 3.**
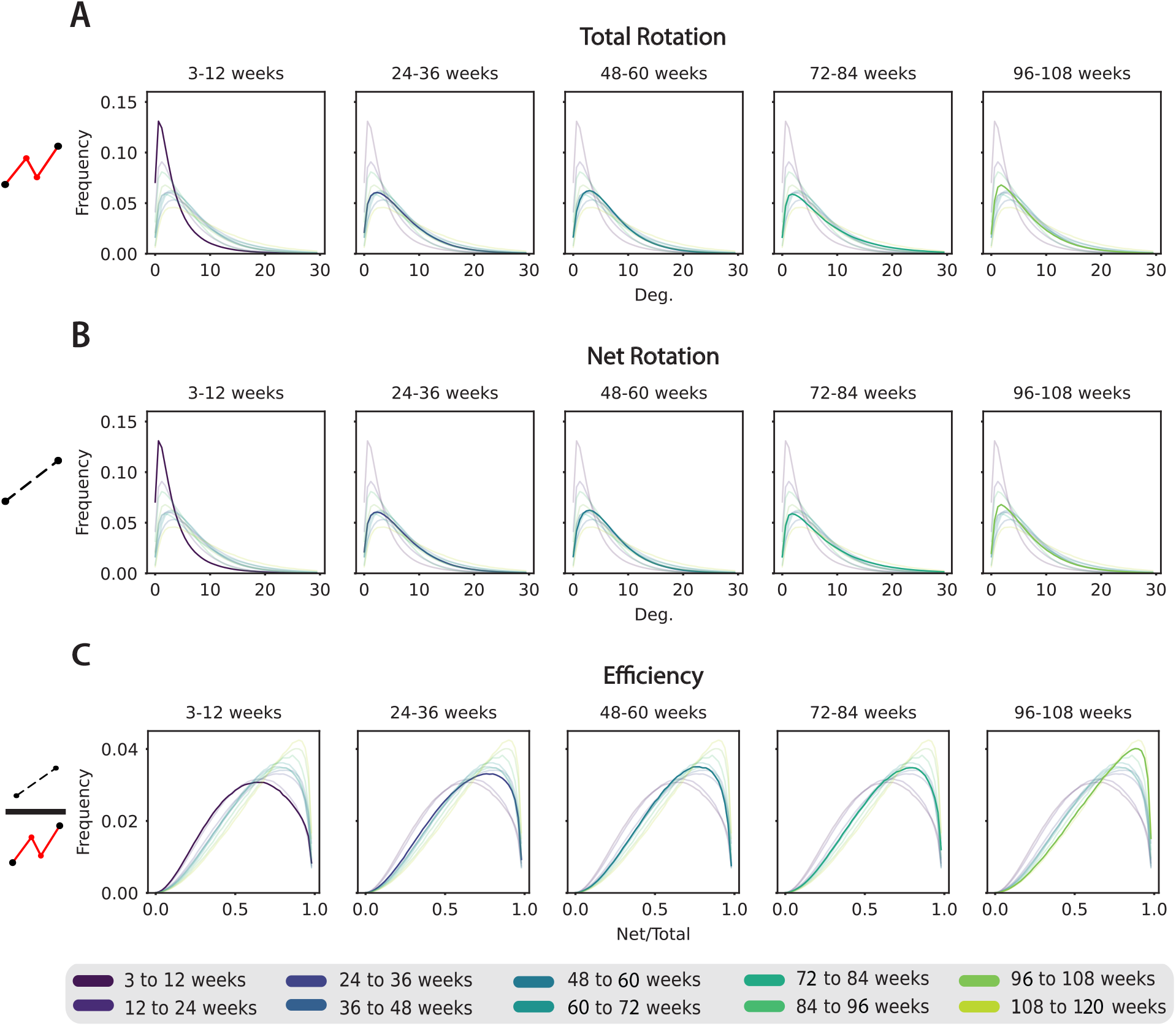
Distributions of head rotation across development. Data are binned by age, with each color representing a 12-week age bin. The panels in each row show the same set of distributions, with a different age group highlighted to enable comparison. **(a) Distributions of total rotation**. The distributions spread progressively outward up to approximately 48 weeks, indicating that both the frequency and magnitude of head rotations increase over the first year of life. **(b) Distributions of net rotation**. As in (A), the distributions develop heavier tails over the first year of life. **(c) Distributions of efficiency**. The peak of the distribution shifts rightward with age, indicating increasingly efficient head rotations across development.

**Figure 4.**
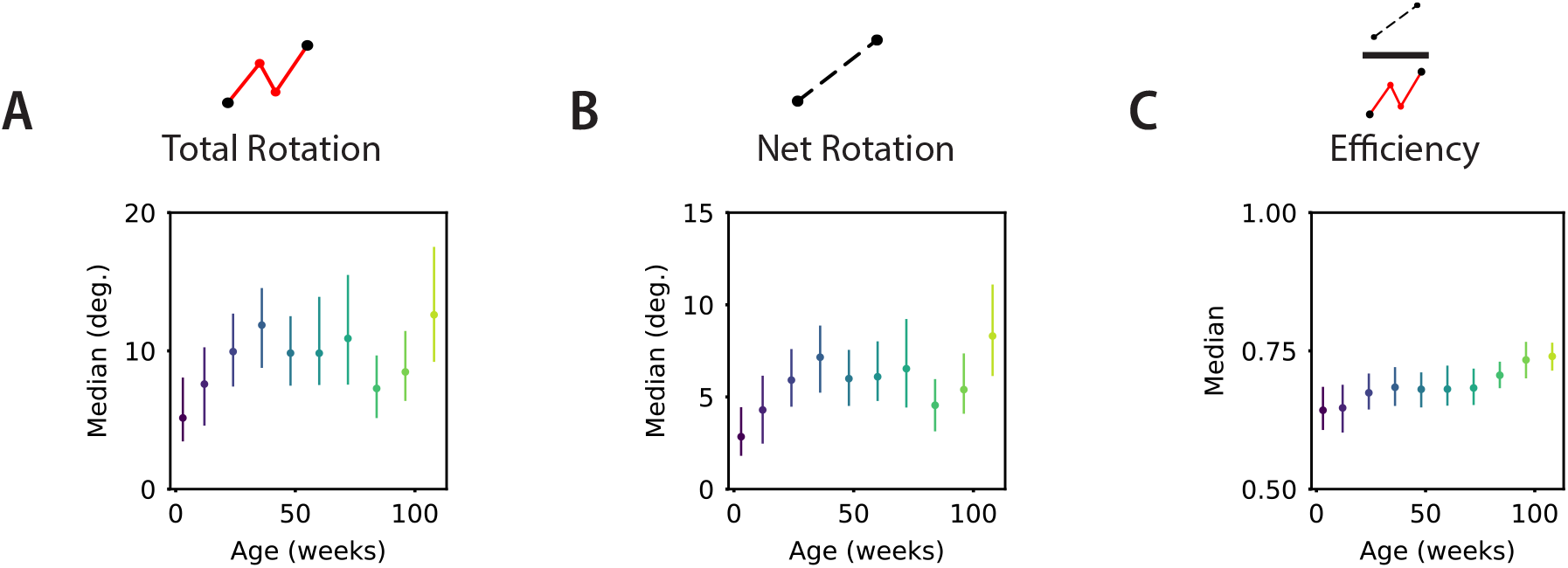
Medians of total rotation, net rotation, and efficiency. **(a)** The median total head-rotation linearly increases during the first 48 weeks of life. **(b)** Consistent with (A), net head-rotation linearly increases during the first 48 weeks. **(c)** Efficiency increases throughout the first 2 years of life, although appearing to plateau from 48 to 72 weeks.

### 2.2 Head rotation efficiency increases over the first two years of life

As infants develop, their coordination improves. We sought to quantify the improvement in head movement coordination in these data by measuring the efficiency of head rotations across development. Across the full age range, rotational efficiency was higher in older infants (*β* = 0.001/week, *SE <* 0.001, *z* = 10.556, *p <* 0.001). Figure 3c shows the distribution of rotational efficiency with age, and Figure 4c shows the median efficiency as a function of age. The increase is particularly evident during 3 to 48 weeks and 84 to 120 weeks.

## 3 Discussion

Our approach measures head rotation as it naturally occurs within the postural and behavioral contexts of daily life. Because postural development and head control are deeply intertwined (Adolph and Franchak, 2017; Bril and Ledebt, 1998), i.e. the postures available to an infant determine the movements the head can and does make, age-related differences in our measures reflect the full system of motor, postural, and attentional changes, not any single component in isolation. This is by design: our goal was to characterize the head movements that actually occur across development rather than to isolate a specific mechanism. However, this means that when we observe, for example, greater rotation magnitudes in older infants, we cannot separate growing neuromuscular ability to direct the head from the fact that older infants are upright and engaged in activities that call for large re-orientations.

The central finding of this work is that head-rotation magnitude and rotational efficiency show distinct age-related developmental patterns during early life. Head-rotation magnitude increased primarily over the first year, suggesting that infants produce larger head movements as strength, posture, mobility, and opportunities for active exploration increase. Rotational efficiency increased across the full age range sampled, through 29 months, indicating that head movements become progressively more directionally coherent with age. These findings suggest that development involves not only changes in how much infants move their heads, but also changes in how those movements are organized. By measuring head rotations during everyday activity at home, this work characterizes head control in the natural contexts in which infants explore objects, interact socially, and move through their environments.

The magnitude of head rotation rises sharply over the first year of life. This early increase likely reflects both expanding motor capacity and the growing use of head turns as a primary means of redirecting visual attention and sampling the environment. After the first year, the median magnitude of rotation fluctuates rather than continuing to rise, and rotational efficiency temporarily plateaus before resuming its increase around 72 weeks. The onset of this fluctuation coincides broadly with the onset of independent walking, which places new demands on head control; head rotations are being incorporated into a broader, more complex system of visual exploration that includes more whole body movement as infants learn to walk. While laboratory studies have documented that motor transitions, like walking, require reorganization of postural and head control (Assaiante and Amblard, 1993; Hadders-Algra et al., 1996; Ledebt and Bril, 2000), this sampling of head movements continuously across development reveals the temporal structure of that reorganization in the context of daily life.

With age, infants’ head rotations became more efficient: a greater proportion of total head movement produced coherent changes in orientation. More efficient rotations should yield more direct shifts in the visual field as infants sample people, objects, and events during everyday activity. This interpretation aligns with head-camera and head-mounted eye-tracking studies showing that infants’ visual input changes with age, posture, locomotor status, and self-generated head movement (Fausey et al., 2016; Kretch et al., 2014; Franchak et al., 2024). Thus, increases in rotational efficiency may help shape the visual information infants generate during natural exploration.

Several limitations qualify these interpretations. Our rotation estimates rely on a visual-odometry pipeline whose performance may vary with scene texture, lighting, motion blur, and independently moving objects, each of which is common in naturalistic egocentric video. If recording conditions differ systematically across ages, these factors could introduce biases into the developmental trajectory. Translation was excluded because monocular estimates are scale-ambiguous, so the analysis cannot address how locomotion couples with head rotation, a gap that is particularly relevant to interpreting the reorganization around walking onset. Finally, the approach does not capture behavioral context: we observe changes in head movement but cannot determine what the infant was doing, attending to, or interacting with at the time.

Future work should address these gaps by combining head-mounted video with complementary measures such as eye tracking, body pose estimation, and activity annotations. Integrating head rotation with richer behavioral data would make it possible to test the motor reorganization and attentional control interpretations more directly. Despite these limitations, our findings offer a new perspective on the development of head control. Two key components of head control, head-rotation magnitude and rotational efficiency, emerge on different timelines, shaping the infant visual experience. Understanding when and how each capacity emerges is essential for characterizing the visual world that supports early learning.

## 4 Materials and Methods

### 4.1 Data collection

These data were collected according to principles of the Declaration of Helsinki under IRB-approved protocols. The procedures for recruiting, consenting, data collection, and data management were reviewed and approved by the Institutional Review Board at Indiana University, and conducted under Protocol no. 1505862312. Families were recruited from an opt-in data base of families interested in participating in research and from outreach events in the community for families of young children. Parents were contacted through mail or email about the methods and goal of this study. If they were interested in participating, an appointment was arranged for training in the use of the head camera and consent was obtained.

Approximately 383 hours of video (*≈* 41 million frames) were collected from 88 infant participants. Egocentric video data were collected using a lightweight, battery-powered Looxcie2 camera secured to the head using a snug-fitting hat. Parents were instructed in the use of the recording system and provided with two hat–camera units. They were asked to orient the camera to the center of the infant ‘s first-person visual field and to record between 4 and 6 hours of video during the infant ‘s waking hours, distributed across naturalistic daily activities in the home environment. These recordings are described in greater detail in Fausey et al. (2016)

### 4.2 Estimating head rotation

Frame-to-frame camera rotations were estimated from the egocentric video using a visual odometry algorithm that segments rigid motion in the scene to isolate camera motion (Yang and Ramanan, 2021). For each consecutive pair of video frames, the algorithm outputs a 3D rotation matrix representing the relative change in camera orientation from frame *t* to frame *t* + 1. Because the camera was securely mounted to the head, estimated camera rotations were attributed to head rotation relative to the environment.

Analyses of these extracted rotations were performed over sliding temporal windows of *N* = 15 frames (corresponding to 0.5 s at 30 Hz) with a stride of one frame. This time window was chosen to approximate the average duration of a head movement during a gaze shift in adults (Fang et al., 2015).

### 4.3 Filtering rotation estimates

To reduce high-frequency noise in the estimated head rotations, relative rotations were filtered prior to analysis using a multi-stage procedure. Rotations were first converted from rotation matrices to a 3D rotation-vector representation and treated as time series sampled at 30 Hz.

Outlier rotations that deviated strongly from the local distribution were then identified based on rotation magnitude using a robust, median-based criterion: rotations exceeding 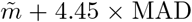, where 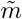 is the median rotation magnitude and 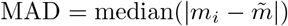 is the median absolute deviation. These outlier samples were replaced via linear interpolation from neighboring rotations to preserve temporal continuity.

The resulting rotation-vector time series was subsequently low-pass filtered using a second-order Butterworth filter with a cutoff frequency of 5 Hz (normalized cutoff *f*_*c*_*/*(*f*_*s*_*/*2) = 5*/*15 = 0.33, where *f*_*s*_ = 30 Hz). This cutoff was chosen to suppress high-frequency noise while preserving behaviorally meaningful head-movement dynamics. Filtering was applied in a zero-phase format to avoid introducing temporal distortions. Finally, the filtered rotation vectors were converted back to rotation matrices for all subsequent analyses.

### 4.4 Verifying head rotation estimates in the Looxcie camera

We designed a controlled moving camera experiment to test the accuracy of the rotation estimation and filtering protocol. We created a large protractor in the lab and placed the camera at its center. During the experiment, we rotated the camera around the yaw axis. We found the algorithm ‘s yaw estimates followed ground truth values closely. The mean absolute position error was 6.27° with a standard deviation of 4.69°. For velocity, the mean absolute error was 0.070°/frame with a standard deviation of 0.056°/frame. The video was recorded at 30 frames per second.

Our analyses measure net rotation (rotation between the 1st and 15th frame) and total rotation (accumulated rotation over 15 frames). Based on our measurements of the per frame error in these controlled experiments, we believe that the error in our velocity measurements over each .5 second period should be limited to relatively small errors, at an average of 2.1°/s.

### 4.5 Total rotation distance

The magnitude of each individual rotation between pairs of frames was extracted using the `scipy.spatial.transform.Rotation` module. Each rotation matrix **R**_*i*_ was converted to an axis–angle representation, and the corresponding rotation angle *θ*_*i*_ was used as the magnitude of rotation. Total rotation or distance within a window was defined as the sum of individual rotation magnitudes, irrespective of their directions:

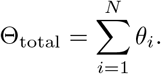

This measure captures the cumulative amount of rotational motion within the window, independent of directional consistency.

### 4.6 Net rotation

The equivalent direct displacement between the beginning and end of the window was determined by multiplying the individual rotation matrices:

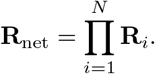

The magnitude of this net rotation, denoted Θ_net_, was then extracted. This quantity estimates the shortest rotation from the starting to ending position for each window, effectively removing directional cancellations between successive rotations.

### 4.7 Rotational efficiency

Rotational efficiency was defined as the ratio of net rotation to total rotation:

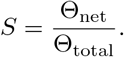

These efficiency values lie in the interval [0, 1], with values near 1 indicating highly consistent, unidirectional rotations, and values near 0 indicating jittery motion during the 0.5 s window. This normalization reveals the directional consistency in the magnitude of the movement.

### 4.8 Statistical analyses

To assess age-related changes in head rotation, we used linear mixed-effects models estimated with restricted maximum likelihood (REML) using the `statsmodels` package (Seabold and Perktold, 2010). Following Petroff et al. (2025), each recording was segmented into non-overlapping 5-minute intervals, and the median value of the measure of interest (total rotation, net rotation, or rotational efficiency) was computed within each interval to serve as the unit of analysis. Age in weeks was included as a fixed effect and infant identity as a random intercept to account for repeated observations within participants. To test whether the age-related trend in rotation magnitude differed across developmental periods, separate models were also fit for infants under 48 weeks and infants 48 weeks and older. Rotational efficiency was modeled across the full age range.

## References

Adolph, K. E. and Franchak, J. M. (2017). The development of motor behavior. Wiley Interdisciplinary Reviews. Cognitive Science, 8(1-2).

Adolph, K. E. and Hoch, J. E. (2019). Motor development: Embodied, embedded, enculturated, and enabling. Annual Review of Psychology, 70(Volume 70, 2019):141–164.

Assaiante, C. and Amblard, B. (1993). Ontogenesis of head stabilization in space during locomotion in children: Influence of visual cues. Experimental Brain Research, 93(3):499–515.

Bertenthal, B. and von Hofsten, C. (1998). Eye, head and trunk control: The foundation for manual development. Neuroscience & Biobehavioral Reviews, 22(4):515–520.

Borjon, J. I., Abney, D. H., Yu, C., and Smith, L. B. (2021). Head and eyes: Looking behavior in 12-to 24-month-old infants. Journal of Vision, 21(8):18.

Bril, B. and Ledebt, A. (1998). Head coordination as a means to assist sensory integration in learning to walk. Neuroscience and Biobehavioral Reviews, 22(4):555–563.

Claxton, L. J., Strasser, J. M., Leung, E. J., Ryu, J. H., and O‘Brien, K. M. (2014). Sitting infants alter the magnitude and structure of postural sway when performing a manual goal-directed task. Developmental Psychobiology, 56(6):1416–1422.

Fang, Y., Nakashima, R., Matsumiya, K., Kuriki, I., and Shioiri, S. (2015). Eye-head coordination for visual cognitive processing. PLOS ONE, 10(3):1–17.

Fausey, C. M., Jayaraman, S., and Smith, L. B. (2016). From faces to hands: Changing visual input in the first two years. Cognition, 152:101–107.

Franchak, J. M. (2020). The ecology of infants’ perceptual-motor exploration. Current Opinion in Psychology, 32:110–114.

Franchak, J. M., Kadooka, K., and Fausey, C. M. (2024). Longitudinal relations between independent walking, body position, and object experiences in home life. Developmental Psychology, 60(2):228–242.

Friedman, A. H., Watamura, S. E., and Robertson, S. S. (2005). Movement–attention coupling in infancy and attention problems in childhood. Developmental Medicine & Child Neurology, 47(10):660–665.

Hadders-Algra, M., Brogren, E., and Forssberg, H. (1996). Ontogeny of postural adjustments during sitting in infancy: Variation, selection and modulation. The Journal of Physiology, 493(Pt 1):273–288.

Klingberg, T., Forssberg, H., and Westerberg, H. (2002). Training of Working Memory in Children With ADHD. Journal of Clinical and Experimental Neuropsychology, 24(6):781–791.

Kretch, K. S., Franchak, J. M., and Adolph, K. E. (2014). Crawling and Walking Infants See the World Differently. Child Development, 85(4):1503–1518.

Ledebt, A. and Bril, B. (2000). Acquisition of upper body stability during walking in toddlers. Developmental Psychobiology, 36(4):311–324.

Mendez, A. H., Yu, C., and Smith, L. B. (2024). Controlling the input: How one-year-old infants sustain visual attention. Developmental Science, 27(2):e13445.

Petroff, Z. J., Jayaraman, S., Smith, L. B., Candy, T. R., and Bonnen, K. (2025). The world through infant eyes: Evidence for the early emergence of the cardinal orientation bias. Proceedings of the National Academy of Sciences, 122(16):e2421277122.

Seabold, S. and Perktold, J. (2010). statsmodels: Econometric and statistical modeling with Python. In Proceedings of the 9th Python in Science Conference, pages 92–96.

Smith, L. B. (2013). It’s all connected: Pathways in visual object recognition and early noun learning. American Psychologist, 68(8):618–629.

Smith, L. B., Yu, C., Yoshida, H., and Fausey, C. M. (2015). Contributions of Head-Mounted Cameras to Studying the Visual Environments of Infants and Young Children. Journal of Cognition and Development, 16(3):407–419.

Yang, G. and Ramanan, D. (2021). Learning to Segment Rigid Motions from Two Frames.

Yu, C. and Smith, L. B. (2012). Embodied attention and word learning by toddlers. Cognition, 125(2):244–262.

